# Skillful prediction of tropical Pacific fisheries provided by Atlantic Niños

**DOI:** 10.1101/2021.02.17.431587

**Authors:** Iñigo Gómara, Belén Rodríguez-Fonseca, Elsa Mohino, Teresa Losada, Irene Polo, Marta Coll

## Abstract

Tropical Pacific upwelling-dependent ecosystems are the most productive and variable worldwide, mainly due to the influence of El Niño Southern Oscillation (ENSO). ENSO can be forecasted seasons ahead thanks to assorted climate precursors (local-Pacific processes, pantropical interactions). However, owing to observational data scarcity and bias-related issues in earth system models, little is known about the importance of these precursors for marine ecosystem prediction. With recently released reanalysis-nudged global marine ecosystem simulations, these constraints can be sidestepped, allowing full examination of tropical Pacific ecosystem predictability. By complementing historical fishing records with marine ecosystem model data, we show herein that equatorial Atlantic Sea Surface Temperatures (SSTs) constitute a superlative predictability source for tropical Pacific marine yields, which can be forecasted over large-scale areas up to 2 years in advance. A detailed physical-biological mechanism is proposed whereby Atlantic SSTs modulate upwelling of nutrient-rich waters in the tropical Pacific, leading to a bottom-up propagation of the climate-related signal across the marine food web. Our results represent historical and near-future climate conditions and provide a useful springboard for implementing a marine ecosystem prediction system in the tropical Pacific.

## 1. Introduction

Before COVID-19, global fisheries supplied society with ~96 million tons of fish. With 39 million people engaged in the primary sector and 4.6 million fishing vessels in operation, the fisheries industry exported USD 164 billion annually [1].

Despite the enormous anthropogenic pressure these numbers reveal, year-to-year variations in marine resources from certain oceanic regions are still largely determined by Earth’s natural variability [2, 3]. This is the case of the tropical Pacific, one of the most biodiverse and productive marine areas worldwide, where ENSO exerts a paramount influence [4–6]. During El Niño events, weakened trade winds allow the penetration of warm waters over the central and eastern tropical Pacific, leading to suppressed equatorial and coastal upwelling of nutrient-rich waters. During La Niña these conditions reverse, inducing colder SSTs, enhanced near-surface nutrient supply and a subsequent response of marine life [7–11].

ENSO is currently the most predictable oceanic-atmospheric variability pattern at seasonal timescales (6-12 months ahead [12]). Relevant ENSO precursors encompass local-Pacific processes [13, 14] and pantropical climate interactions [15]. Recent evidence suggests the latter are more important than previously thought in shaping ENSO evolution, intensity and structure [16–21]. Pantropical interactions are generated by recurrent SST anomaly patterns from the tropical Atlantic and Indian oceans (e.g., equatorial and North Tropical Atlantic [22–24], Indian Ocean Dipole [25] and Indian Ocean Basin Mode [26]). SST fluctuations are sufficiently energetic to alter the Walker circulation and the surface wind regimes in the tropical Pacific [16–19, 27, 28]. However, ENSO dynamics associated with remote SST forcing are disparate.

Boreal spring North Tropical Atlantic (NTA) and autumn Indian ocean SST forcing mainly induce advective processes over the western tropical Pacific [19, 29, 30]. Distinctively, equatorial Atlantic boreal summer anomalous SSTs (a.k.a., Atlantic Niños/Niñas) can excite eastward-propagating oceanic Kelvin waves in the eastern Pacific [31, 32], thereby altering the thermocline and the upwelling of ocean bottom waters. The settled environmental conditions are in all cases conducive to ENSO events in the ensuing seasons, but their impact on equatorial and coastal upwelling is different [15, 23]. Thus, it is important to assess their relevance for marine ecosystem prediction, a fracture that has remained uncertain for several reasons.

Scarce, short or too local historical biological records often hinder the identification of robust statistical links between climate predictors and yield responses [33], especially when considering vast areas like the tropical Pacific. Dynamical prediction by means of effectively coupled climate, biogeochemical and ecosystem models has been proven successful over multiple coastal and open sea regions [34–36]. However, while most state-of-the-art freely-evolving driving climate models are good at predicting ENSO [12, 37], they fail to reproduce observed pantropical interactions due to large SST systematic errors [15, 38, 39]. Thus, critical remote predictability sources of tropical Pacific ecosystems may be overlooked (e.g., Atlantic Niños).

Alternatively, high-resolution ecosystem models nudged to earth system reanalyses, like those recently released by the Fisheries and Marine Ecosystem Model Intercomparison Project (FishMIP [40]), enable to sidestep large SST model biases, fully consider ENSO precursors and implement rigorous model-based statistical ecosystem prediction. Released fished and non-fished FishMIP historical simulations not only permit the effective isolation of climate-driven ecosystem responses, but also characterize the top-down control of this anthropogenic forcing.

Given the above, we evaluate herein the potential predictability of tropical Pacific marine ecosystems and fisheries by assessing the precursory role of global SSTs. To this end, we utilized atmospheric-oceanic reanalysis data, historical catch records and FishMIP simulations encompassing global marine ecosystems.

## 2. Materials and Methods

### 2.1 Observational datasets and reanalyses

Annual catch data from FAO major fishing areas *Eastern Central Pacific* (#77) and *Southeastern Pacific* (#87) for the period 1950-2014 were retrieved from the Sea Around Us Project (SAUP) database. These are a combination of official reported data from FAO FishStat database (http://www.fao.org/fishery/statistics/en) and reconstructed estimates of unreported data [41]. To retain interannual variability and minimize the potential impact of external factors (e.g., fishing effort, climate change impact) on historical catch data, a high-pass Butterworth filter with 11-year cut-off period was applied to the time series.

Observational 1° longitude-latitude SST data were retrieved from the Hadley Centre Sea Ice and Sea Surface Temperature data set (HadISST [42]).

Analogous reanalysis data were obtained from the GFDL Carbon Ocean Biogeochemistry and Lower Trophics run (COBALT [43]). Physical-biological parameters from GFDL reanalysis and additional datasets are fully described in *Supplementary Material* Table S1. The COBALT ocean-ice simulation, performed with the Modular Ocean Model (MOM) v4p1 [44], has a 1° horizontal resolution (1/3° along the equator) and 50 vertical layers with 10 m increase in the top 200 m. The simulation is forced by air-sea fluxes from the CORE-II data set [45], a blend of the National Centers for Environmental Prediction (NCEP) atmospheric reanalysis [46] and satellite data. Full information of the GFDL COBALT calibration/validation against physical-biological observations (e.g., mixed layer depth, nutrient concentration, phytoplankton/zooplankton biomass, carbon/energy fluxes) is available in Stock et al. [43].

Consistent with CORE-II forcing on GFDL COBALT, NCEP reanalysis was selected to analyze atmospheric circulation dynamics and variability in this work (cf. *Supplementary Material* Table S1).

### 2.2 Fisheries and Marine Ecosystem Model simulations

Model simulations from the Fish-MIP v1 intercomparison protocol [40] were analyzed. Selected available models were EcoOcean, BOATS and Macroecological [47].

EcoOcean is a complex spatial-temporal global ecosystem model that comprehensively simulates interactions within all trophic levels and species groups using the Ecospace [48] spatial-temporal model at its core. The depth dimension is implicitly considered in the model by preferent habitat patterns and food-web interactions. EcoOcean v1 uses SAUP effort data [49] spatially distributed across large marine ecosystem regions as fishing forcing. The model is able to replicate historical catch successfully on a global scale. A full description of EcoOcean is available in Christensen et al. [50].

BOATS is a global ecological model that determines size spectra of fish biomass as a function of net primary production and local temperature. Biomass production in higher tropic levels is limited by available photosynthetic energy, temperature-dependent growth rates and trophic efficiency scaling. The model enables a realistic representation of biological and ecological processes, allowing evaluation of the links between climate, biogeochemistry and upper trophic dynamics. BOATS employs a bioeconomic approach based on SAUP catch price data to determine spatial-temporal changes in fishing efforts. Full information on the model architecture is available in Carozza et al. [51].

Macroecological is a static-equilibrium model that relies on predator-prey mass ratios, transfer efficiency and metabolic demands dependent on body mass and temperature to predict size and distribution of marine consumers (as a whole) at any given time and location. It has a single vertical (surface-integrated) layer and movement of fish is disregarded. Extended information on the model is available in Jennings and Collingridge [52].

Six different simulations were analyzed in this study. These include fished and unfished runs with and without diazotrophs dynamics. All historical runs (1971-2004) are forced by GFDL COBALT reanalysis, whereas an EcoOcean 2021-2054 projection is driven by IPSL-CM5A-LR RCP8.5 [53]. In the latter, fishing forcing is kept constant at 2005 rates [40]. Although GFDL-ESM2M was also available as driving forcing for 2021-2054, it was eventually disregarded due to its inability to simulate a connection reminiscent of the observed Atlantic-Pacific teleconnection subject of this study [54]. Complete information on simulation settings, acronyms and data sources are available in *Supplementary Material* Table S2. All common FishMIP model output variables considered are detailed in *Supplementary Material* Table S1 and are publicly available through the FishMIP global model repository [47]. The most detailed model (EcoOcean) forced by historical fisheries effort is used as reference throughout the manuscript (hereafter EcoG-Fis), as it provides the closest possible approximation to reality.

### 2.3 Physical-biological indices and statistical methods

Monthly anomalies of physical-biological variables were calculated by removing their total period average to the time series of each grid point. Seasonal (4-month) anomalies were computed as aggregated monthly anomalies, with linear trends effectively removed to minimize the impact of anthropogenic climate change [55].

Physical-biological indices were determined by spatially averaging seasonal anomalies over regions of interest, e.g., Niño3 [150°W-90°W, 5°S-5°N] and Atl3 [20°W-0°E, 3°S-3°N]. All indices were normalized by dividing the time series by their standard deviation.

Whereas Atl3 was computed for SST on boreal summer (July to September; JJAS) to represent Atlantic Niño variability during its peak intensity [22] (a.k.a. Atlantic Equatorial Mode), Niño3 and global seasonal anomalies of physical-biological variables (cf. *Supplementary Material* Table S1) were calculated as rolling 4-month averages between monthly lags −8 to +24, where lag 0 corresponds to JJAS, lag1 (−1) to JASO (MJJA) of the same calendar year and so forth (back).

Statistical significance for correlation outputs (Figs. 1, 3 and 4; 95% confidence interval) was evaluated using a Student-t test that accounts for the autocorrelation of the time series and the calculation of effective degrees of freedom [56].

**Fig. 1.**
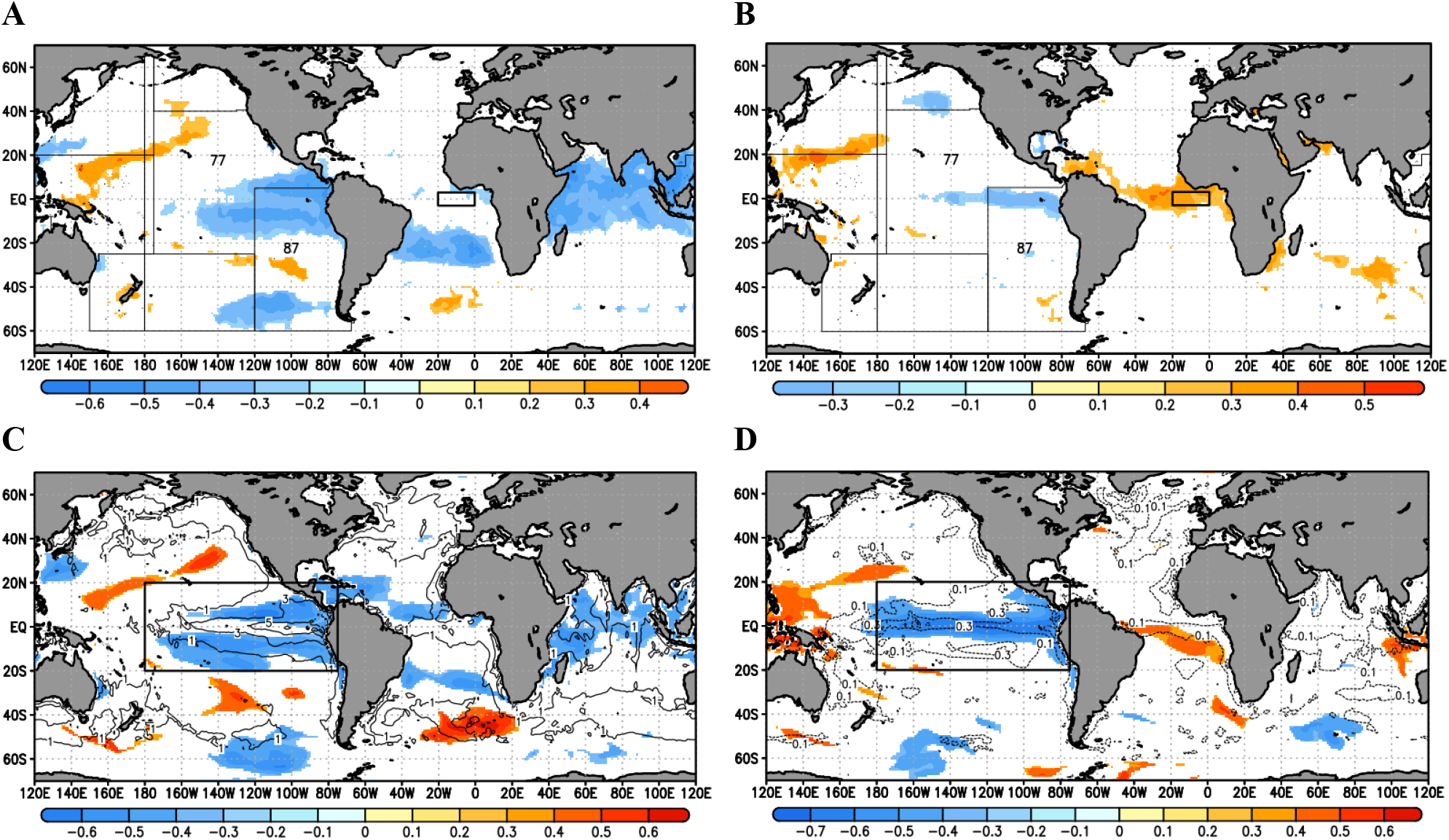
Links between Pacific total catch and global SSTs. (*A*) Correlation between interannual Sea Around Us Project historical catch (FAO aggregated 77 & 87 areas) anomalies and concomitant SST (HadISST) anomalies (shadings; p-value<0.05). Period 1950-2014. Box for Atl3 SST index calculation in black rectangle. (*B*) Same as but for correlation with previous calendar year SST anomalies. (*C*) Correlation between interannual tropical Pacific averaged (black rectangle) simulated historical total catch anomalies from EcoOcean model (EcoG-Fis simulation) and concomitant SST anomalies from GFDL-COBALT (shadings; p-value<0.05). Period 1971-2004. Annual averaged carbon biomass density of commercial species (*bcom*) from EcoOcean model in contours. (*D*) Same as (*B*) but for correlation with previous calendar year SST anomalies (shadings) and standard deviation of annual *bcom* (contours).

### 2.4 S4CAST model

The Sea Surface Temperature-based Statistical Seasonal forecast model (S4CAST) is a novel statistical seasonal model that utilizes SST data (predictor) to forecast the response of any given impact variable (predictand). To this end, the model performs a Maximum Covariance Analysis (MCA [57]) between a predictor Y field (SST anomalies) and a predictand Z field for any given region, time period and lag.

MCA is based on Single Value Decomposition (SVD) applied to the non-square covariance matrix (C) of both Y and Z fields. The method calculates linear combinations of the time series of Y and Z (a.k.a. expansion coefficients U and V, respectively) to maximize C.

A leave-one-out cross-validated hindcast based on MCA outputs is then performed [58]. For this purpose, predictor/predictand data combinations from all years except the year being predicted at each training stage are selected for model training and validation [58, 59]. A Monte Carlo test with 1,000 random permutations is applied to MCA (U vs. anomalous field correlations) and cross-validation of skill scores for hypothesis testing (95% confidence interval).

In this work, Atl3 JJAS SST anomalies were set as predictor, while tropical Pacific [150°W-75°W, 20°S-20°N] simulated total catch and total system biomass anomalies were used as predictands for different monthly lagged periods (0 to 24; EcoG-Fis simulation). The Pearson correlation coefficient was selected as a qualitative indicator for model skill (sign of response). Further details on the S4CAST statistical model framework are available in Suárez-Moreno and Rodríguez-Fonseca [60].

## 3. Results

### 3.1 Global SST forcing on tropical Pacific fisheries

We find year-to-year variations of historical global SSTs and eastern Pacific total catch records from the Sea Around Us Project (SAUP) database (FAO areas 77 & 87), robustly correlated for the period 1950-2014. The years when observed fish catches are higher than usual synchronize with well-developed La Niña-like SST anomalies over the tropical Pacific and elsewhere [11, 61] (Fig. 1*A*). Lagged SST anomalies from the previous calendar year depict a preceding Atlantic Niño signal, together with a developing La Niña over the Pacific (Fig. 1*B*).

Likewise, we consider EcoOcean model historical simulation in a fishing forcing scenario (EcoG-Fis) for the same analysis. We first characterize the tropical Pacific in terms of biomass density of commercial species. As expected, this is the most productive and fluctuating region worldwide in the model (Fig. 1*C-D*). Despite the slightly smaller area and shorter period considered (FishMIP coverage; 1971-2004), results very similar to observations are obtained from the model (Fig. 1*C-D*). Apart from equatorial Atlantic SST anomalies and intrinsic 2-7-yr Niña-Niño intermittency, no other significant relations with tropical SST anomalies (e.g., NTA, Indian) are found, either in observations or in simulations during the years preceding annual-averaged total catch anomalies (Fig. 1 and *Supplementary Material* Fig. S1).

Correlation between the Atlantic Niño index (Atl3 JJAS SST; hereafter Atl3-Had) and the tropical Pacific total catch corroborates the lagged 1-yr Atlantic-Pacific relationship in the model and observations (*Supplementary Material* Fig. S2). As expected, this relationship substantially weakens and changes sign in subsequent year lags. In agreement with Rodríguez-Fonseca et al. [16], a slightly stronger Atlantic-Pacific connection is found for 1971-2004 compared to 1950-2014.

### 3.2 Statistical prediction of tropical Pacific total catch using equatorial Atlantic SSTs

Owing to constraints in historical fishing records (averaged annual data over vast areas), we focus on high resolution monthly FishMIP modelling outputs for fisheries prediction. Again, we consider the most detailed model forced by historical fisheries efforts (EcoG-Fis), which provides the closest approximation to reality (cf. Supplementary Material Table S2). We then use the Sea Surface temperature-based Statistical Seasonal forecast model (S4CAST [60]) to perform a cross-validated hindcast of tropical Pacific total catch at sea (tc) simulated by the model.

Consistent with Fig. 1, we use as predictor equatorial Atlantic SST anomalies during the peak intensity season (JJAS-lag 0) of Atlantic Niños/Niñas [16, 22, 62] on GFDL-COBALT, the driving ocean reanalysis of FishMIP.

For all monthly lags, Atlantic Niño emerges as the main predictor of tropical Pacific total catch anomalies, which can be hindcasted over large-scale areas and up to two years in advance (Fig. 2 and *Supplementary Material* Fig. S3).

**Fig. 2.**
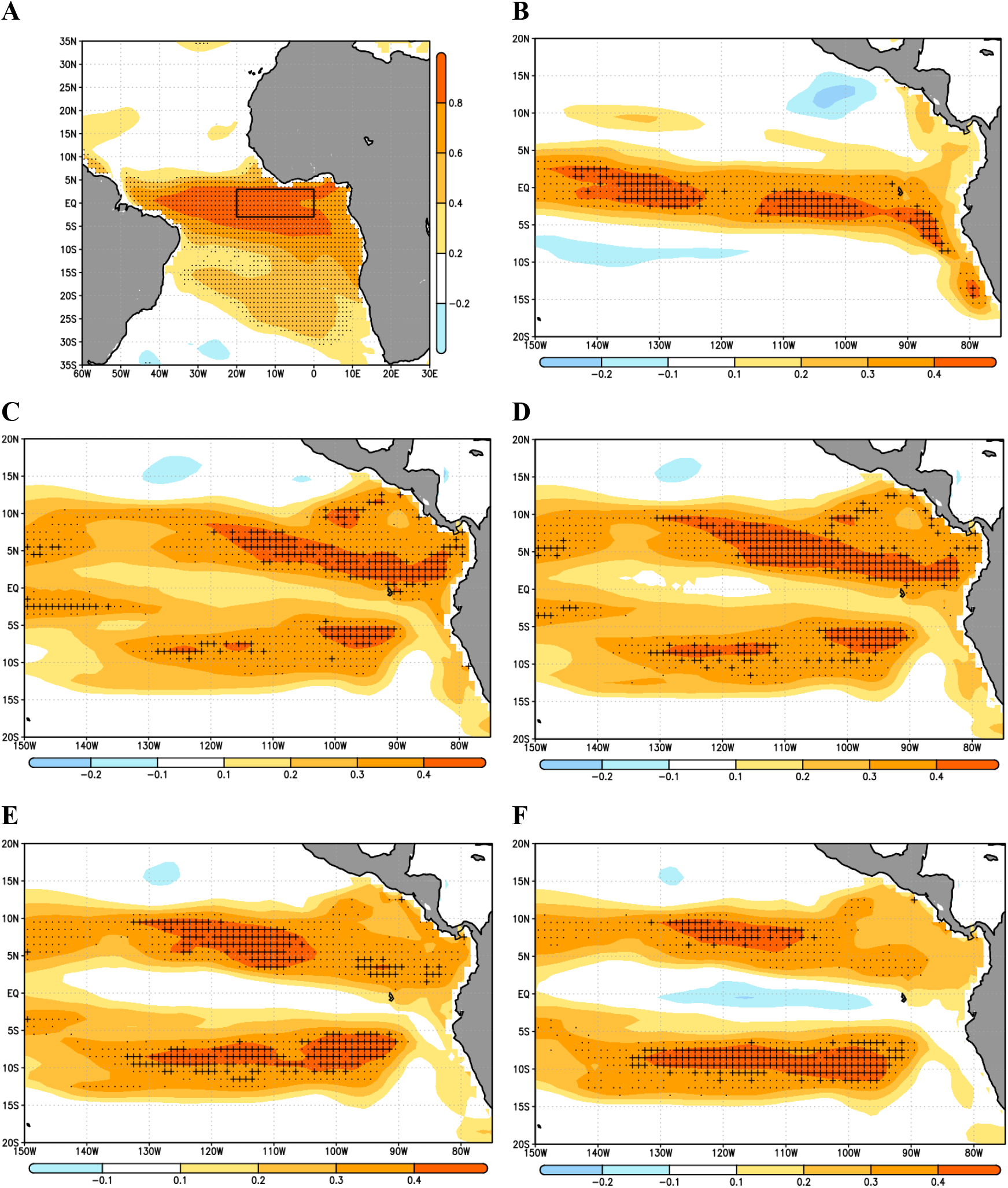
Hindcasts of tropical Pacific total catch based on Atlantic SSTs. Co-variability maps corresponding to the leading Maximum Covariance Analysis (MCA) modes between equatorial Atlantic lag 0 SST anomalies (predictor; Atl3 region [20°W-0°E, 3°S-3°N] - black box) and tropical Pacific total catch (*tc*) at sea at different monthly lags (predictand; [150°W-75°W, 20°S-20°N]). (*A*) Homogeneous map obtained by regression of the expansion coefficient U on tropical Atlantic SST anomalies on monthly lag 0 (JJAS). P-values<0.05 in SST-U correlation in stippling (Monte Carlo test with 1,000 random permutations). (*B*) Heterogeneous map obtained by regression of U on tropical Pacific *tc* anomalies on monthly lag 4 (ONDJ). P-values<0.05 in *tc*-U correlation in stippling. Cross-validated anomaly correlation skill scores calculated from the MCA between hindcasts and observations of *tc* in crosses (95% confidence interval). (*C-F*) Same as (*B*) but for monthly lags 18 (DJFM), 20 (FMAM), 22 (AMJJ) and 24 (JJAS). Period 1971-2004. EcoG-Fis simulation. See *Methods* for further information on MCA.

### 3.3 Atlantic-Pacific physical-biological mechanism

To establish a physical-biological mechanism explaining the results above, we evaluate SST tropical Atlantic (Atl3) impact on Pacific (Niño3) region for different area-averaged physical-biological variables and monthly lags. The simulation selected for this purpose is again EcoG-Fis, although other marine ecosystem models (BOATS, Macroecological) and scenarios are also considered (cf. *Supplementary Material* Tables S1-S2 and Tittensor et al. [40]).

First, we check whether the Atlantic-Pacific physical teleconnection [16, 28] is realistically represented in GFDL-COBALT. This is confirmed in Fig. 3*A* and *Supplementary Material* Fig. S4. As expected, for a warming in the equatorial Atlantic, a strengthening of the Pacific Walker Circulation is observed, together with increased subsidence and anomalous easterlies over the western equatorial Pacific, features that promote subsequent La Niña conditions (*Supplementary Material* Fig. S4). Similarly, previous studies have shown that anomalous easterly winds over the western equatorial Pacific promote the propagation of equatorial upwelling Kelvin waves [31], thus favoring the emergence of nutrients from the seabed.

**Fig. 3.**
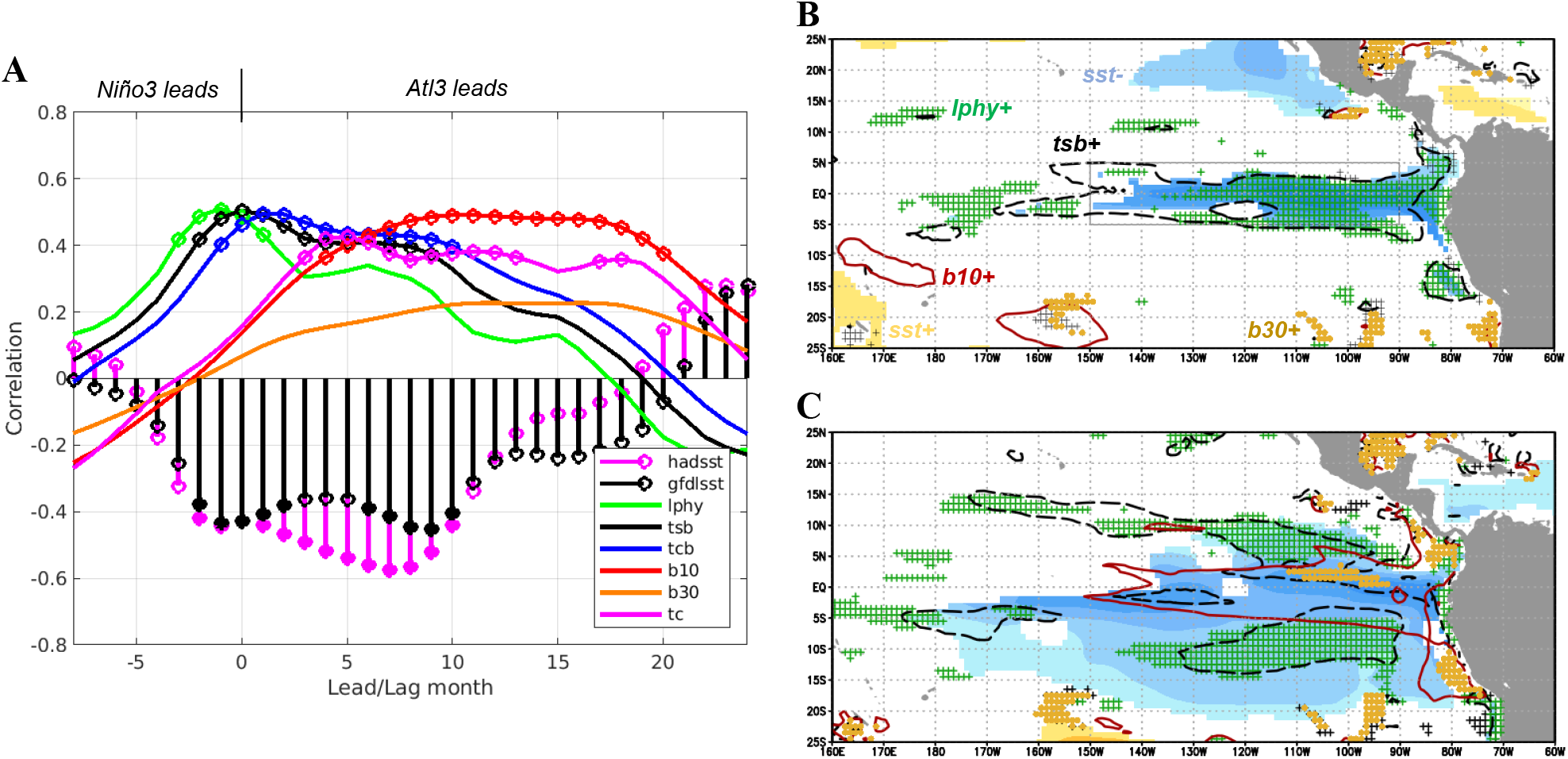
Atlantic-Pacific physical-biological mechanism. (*A*) Correlation between Atl3-GF index and lagged (−6 to 24) Niño3 rolling 4-month anomalies of SST, large phytoplankton production (*lphy)*, total system carbon biomass density (*tsb)*, total consumer carbon biomass density (*tcb*), carbon biomass density of consumers greater than 10 cm (*b10)* and 30 cm (*b30*) and total catch at sea (*tc*). P-values<0.05 in markers (see legend). Regression of tropical Pacific lag 0 SST anomalies (shadings; in K) on Atl3-GF index, and correlation of *lphy* (green crosses), *tsb* (black dashed contours), *b10* (red contours) and *b30* (orange xs) anomalies with the same index. Only p-values<0.05 for positive correlations are shown, except for SST (see text labels). Niño3 region highlighted in thin rectangle. (*C*) Same as (*B*) but for monthly lag 9. Period 1971-2004. GFDL-COBALT and EcoG-Fis simulations.

Lagged-correlations for biological variables in Fig. 3*A* enable characterization of the bottom-up propagation of Atl3-related signal (hereafter Atl3-GF) across the Niño3 region food web. Closely synchronous to the growth of cold nutrient-rich SST anomalies (Fig. 3*A*), large phytoplankton production (*lphy*), total system (*tsb*) and total consumer (*tcb*) carbon biomass densities grow rapidly before monthly lag 0, and decay thereafter at different rates. The different declining rates are due to distinct variable definitions and growth rates of species included in each category. While *lphy* is large-phytoplankton dependent, *tcb* is dominated by zooplankton and *tsb* is a blend of both (cf. *Supplementary Material* Table S1). Consistent with previous studies [63], we find no SST response by small phytoplankton (not shown). Variables negative lag behavior can be partially explained by construction of Atl3 index at its peak season instead of its growing stage (*Supplementary Material* Fig. S5), and the fact that ocean chlorophyll tends to slightly precede ENSO-related SST anomalies due to wind-iron supply dynamics [64].

Results for carbon biomass density of fish consumers greater than 10cm (*b10*) and 30cm (*b30*) depict a bottom-up propagation of the physical forcing over time, peaking ~9-12 months later than large phytoplankton productivity (Fig. 3*A*). By construction of *b10/b30*, where fish species are classified in terms of body size, bottom-up indirect effects of climate anomalies seem to overbalance species-specific direct temperature-recruitment processes [61]. A feasible process for this bottom-up propagation is a recruitment increase of short-lived species caused by enhanced food supply at lower trophic levels [36].

The spatial response of physical-biological variables for lags 0 and 9 is shown in Fig. 3*B-C*. At lag 0, cold SST anomalies and enhanced *lphy/tsb* are present over the Niño3 region (Fig. 3*B*). At lag 9, a mature La Niña is present, together with increased *b10* over the Niño3 area and *lphy/tsb* to the north and south. This northward-southward advection of *lphy* appears related to Ekman transport induced by enhanced trade winds (*Supplementary Material* Fig. S6).

A broader picture of this evolution is given in Fig. 4 in the form of Hovmöller diagrams. Enhanced *lphy/tsb/tcb* tend to overlap with cold SST anomalies, preserving the timing from Fig. 3*A*. Growth starts at small negative lags over the central tropical Pacific, then propagate to the east until lag 3 (Fig. 4*A*) and split thereafter into two branches to the north and south (Fig. 4*B*). The long persistence of *lphy/tsb/tcb* may be caused by enduring presence of nutrients (nitrate and dissolved iron), which unlike SSTs do not interact with the atmosphere [8, 36]. Lastly, whereas *b10* show a similar response to *lphy* ~9 months later, increased *b30* are constrained over the northeast tropical Pacific from lags 14 to 24 (Fig. 4*B*).

**Fig. 4.**
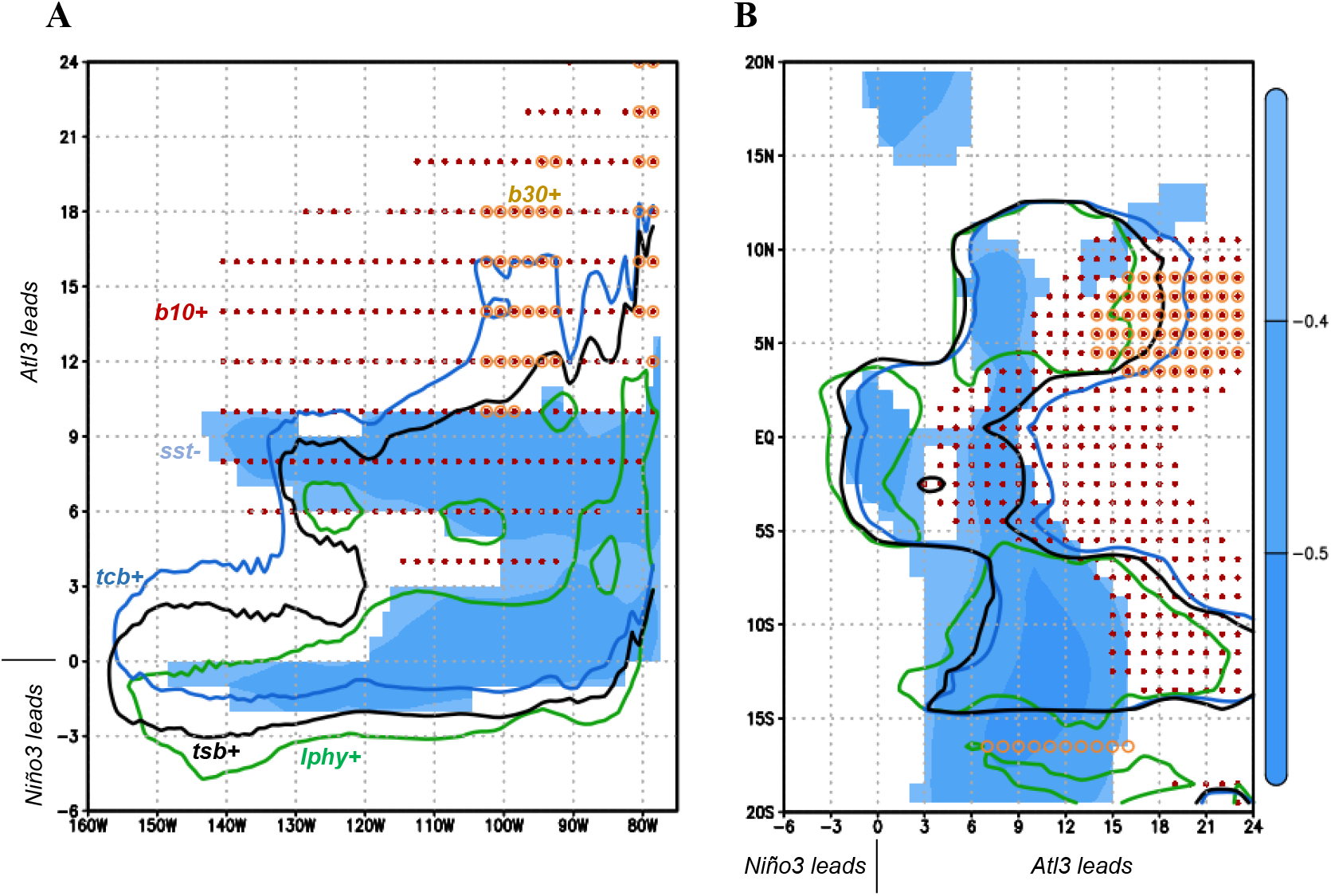
Atlantic-Pacific physical-biological mechanism. Hovmöller diagrams. (*A*) Regression of meridionally averaged [5°S-5°N] rolling 4-month anomalies of SST (shadings; in K) on Atl3-GF index, and correlation of meridionally averaged large phytoplankton production (*lphy;* green contours), total system carbon biomass density (*tsb;* black contours), total consumer carbon biomass density (*tcb;* blue contours) and carbon biomass density of consumers greater than 10 cm (*b10*; red dots) and 30 cm (*b30*; orange circles) with the same index. Only p-values<0.05 for positive correlations are shown, except for SST (see text labels). Lags −6 to 24. (*B*) Same as (*A*) but for zonally averaged anomalies [150°W-75°W]. Period 1971-2004. GFDL-COBALT and EcoG-Fis simulations.

The evolution of physical-biological variables shown in Figs. 3 and 4 are consistent with the shape and progression of total catch predictable regions in Fig. 2 (large crosses). Indeed, if total system carbon biomass density is considered for prediction instead of total catch, final results are very similar but shifted ~9-12 months back in time (compare *Supplementary Material* Figs. S3 and S7). This is consistent with the propagation in time of climate anomalies within the food web (Figs. 3 and 4).

To test the sensitivity of ecosystem response to top-down fishing control and model setup, in *Supplementary Material* Figs. S8-S10 we perform the same analyses on EcoOcean and BOATS fished and unfished model output scenarios. Results indicate that climate-driven bottom-up propagation dominates the response throughout the trophic web, while fishing visibly affects large predators over large-scale areas. The differences observed between models (less persistent and westward displaced anomalies of all-size fish biomasses in BOATS model outputs) may be caused by different internal hypothesis regarding species dynamics, ecosystem functioning and fisheries schemes (cf. *Methods*).

Lastly, a similar analysis is conducted for Macroecological, although it only provides annual model outputs, does not include a fishing scenario, and propagation in time of anomalies through the food chain is unresolved (*Supplementary Material* Fig. S11). As expected, the model returns the same picture as in Fig. 3*B-C* for lagged years 0 and 1, with the difference that consumer response (*tcb*/*b10*/*b30*) to the forcing is no longer lagged in time.

## 4. Summary and discussion

In this work, the predictability of seasonal to multiannual tropical Pacific marine ecosystem resources was investigated on observations (SAUP catch data) and ecosystem model simulations (FishMIP). Evidence indicates that tropical Pacific yield anomalies (e.g., total system biomass, total catch at sea) can be forecasted over large-scale areas, and up to 2 years in advance, by solely considering boreal summer equatorial Atlantic SST anomalies (Atlantic Niños/Niñas) as predictor.

The outstanding skill for predicting Pacific fisheries provided by Atlantic Niños/Niñas can be attributed to the dynamical processes they trigger in the tropical Pacific. These processes involve the alteration of upwelling conditions by anomalous surface winds through the propagation of equatorial Kelvin waves [31, 32]. Contrastingly, other well-known remote SST ENSO precursors are associated with more advective processes over the western Pacific [19, 30] (e.g., NTA, Indian Ocean Dipole), which do not seem to have a substantial impact on high-productive upwelling-dependent marine ecosystems located further to the east in the basin. These results highlight the importance of equatorial Atlantic impact on ENSO [16, 28, 65].

It remains uncertain whether the described biophysical Atlantic-Pacific relation will persist in the future. It is envisaged that under greenhouse warming conditions biomass over the tropical oceans will decline [55], strong eastern ENSO events will become more likely [66] and the Atlantic-Pacific teleconnection (nonstationary in nature [16, 32]) will weaken [54]. Furthermore, future scenarios for fisheries are currently even more uncertain than pre-COVID-19 ones [67].

Although disentangling the interplay of all these factors may require a suite of sensitivity experiments beyond the scope of this study, we provide exploratory results for the period 2021-2054 in *Supplementary Material* Fig. S12 (RCP8.5 fished scenario; cf. *Supplementary Material* Table S2). Overall, results analogous to those obtained for the historical period are found, supporting the applicability of our findings for the decades to come.

Nevertheless, we simply propound a few possible alternative approaches to marine ecosystem modelling. When additional FishMIP simulations become available, a more comprehensible picture may unfold.

## Acknowledgments

This research was funded by the EU H2020 project TRIATLAS (no. 817578), the Universidad Complutense de Madrid project FEI-EU-19-09 and the Spanish Ministry of Economy and Competitiveness project PRE4CAST (CGL2017-86415-R). We thank D. Tittensor (UNEP-WCMC) and Iliusi Vega (PIK-Postdam) for help with FishMIP data extraction. We thank R. Suárez-Moreno (LDEO-Columbia University) and J. Steenbeek (ICM-CSIC) for their cooperation during the progress of this study.

## Data availability

Annual catch data from FAO major fishing areas are available from the Sea Around Us Project (http://www.seaaroundus.org/data/#/fao). Hadley Centre Sea Ice and Sea Surface Temperature data set (HadISST) are available from the Met Office Hadley Centre observations datasets (https://www.metoffice.gov.uk/hadobs/hadisst/). National Centers for Environmental Prediction (NCEP) reanalysis data are available from the NOAA/OAR/ESRL PSL website (https://psl.noaa.gov/). GFDL COBALT and FishMIP simulation data (EcoOcean, BOATS and Macroecological) are accessible through the Potsdam-Institute for Climate Impact Research Earth System Grid Federation (ESGF) data node (https://esg.pik-potsdam.de/search/isimip/).

## Code availability

S4CAST originally published MATLAB® code is open access and available from the Zenodo repository (doi:10.5281/zenodo.15985) in the URL https://zenodo.org/record/15985. The rest of MATLAB scripts used in this analysis can be made available upon request from the corresponding author. Updated S4CAST versions can be requested to the TROPA-UCM research group.

## Notes

### Competing Interest Statement

The authors have declared no competing interest.

