## Supplementary Material for "Skillful prediction of tropical Pacific fisheries provided by Atlantic Niños"

**This Supplementary Material file includes:**

- Tables S1 and S2
- Figures S1 to S12

**Supplementary Tables**

**Table S1.** Description of datasets and variables considered in this study.

| Dataset | Variable (*acronym*) | Units | Notes |
|---|---|---|---|
| | | **Observations and reanalyses** | |
| **SAUP** | Annual Catch | $10^3$ t yr$^{-1}$ | FAO major fishing areas 77 and 87 |
| **HadISST** | Sea Surface Temperature (SST) | K | |
| **GFDL COBALT** | Sea Surface Temperature (SST) | K | |
| | Surface zonal and meridional currents (*uo,vo*) | m s$^{-1}$ | |
| | Small and large phytoplankton production (*sphy* and *lphy*) | mol C m$^{-3}$ s$^{-1}$ | Vertical integrals |
| **NCEP** | Zonal, meridional and vertical winds (*u,v,w*) | m s$^{-1}$ | |
| | Velocity potential (*chi*) | $10^6$ m$^2$ s$^{-1}$ | 0.2 sigma level |
| | Vertical velocity in pressure coordinates (*omega*) | Pa s$^{-1}$ | |
| | | **Marine ecosystem model simulations** | |
| **EcoOcean** | Total system carbon biomass density (*tsb*) | g C m$^{-2}$ | All primary producers and consumers |
| | Total consumer carbon biomass density (*tcb*) | g C m$^{-2}$ | Trophic level>1, invertebrates and vertebrates |
| | Carbon biomass density of consumers greater than 10cm (*b10*) and 30cm (*b30*) | g C m$^{-2}$ | b10 include b30 |
| | Carbon biomass density of commercial species (*bcom*) | g C m$^{-2}$ | Harvested fish greater than 10cm; fishing scenario |
| | Total catch at sea (*tc*) | g wet biomass m$^{-2}$ | Commercial landings, discards, fish and invertebrates; fishing scenario. |
| **BOATS** | Total consumer carbon biomass density (*tcb*) | g C m$^{-2}$ | Only commercial species |
| | Biomass density of small (*bsmall*), medium (*bmed*) and large size (*blarge*) consumers | g C m$^{-2}$ | |
| | Total catch at sea (*tc*) | g wet biomass m$^{-2}$ | Fishes excluding invertebrates; fishing scenario |
| **Macroecological** | Total consumer carbon biomass density (*tcb*) | g C m$^{-2}$ | Annual average. Only commercial species |
| | Carbon biomass density of consumers greater than 10cm (*b10*) and 30cm (*b30*) | g C m$^{-2}$ | Annual average. b10 include b30 |

**Table S2.** Description of available FishMIP simulations analyzed and scenario acronyms.

| Model | Resolution | Diazotrophs | Forcing | Period | Fishing | Acronym |
|---|---|---|---|---|---|---|
| **EcoOcean** | 1ºx1º Monthly | Yes | GFDL COBALT Reanalysis | 1971-2004 | SAUP effort | EcoG-Fis |
| | | | | | No | EcoG |
| | | | IPSL-CM5A-LR RCP 8.5 | 2021-2054 | SAUP effort 2005 | EcoI8.5-Fis |
| **BOATS** | 1ºx1º Monthly | No | GFDL COBALT Reanalysis | 1971-2004 | SAUP catch price | BoaG-Fis |
| | | | | | No | BoaG |
| **Macroecological** | 1ºx1º Yearly | No | GFDL COBALT Reanalysis | 1971-2004 | No | MacG |

### Supplementary Figures

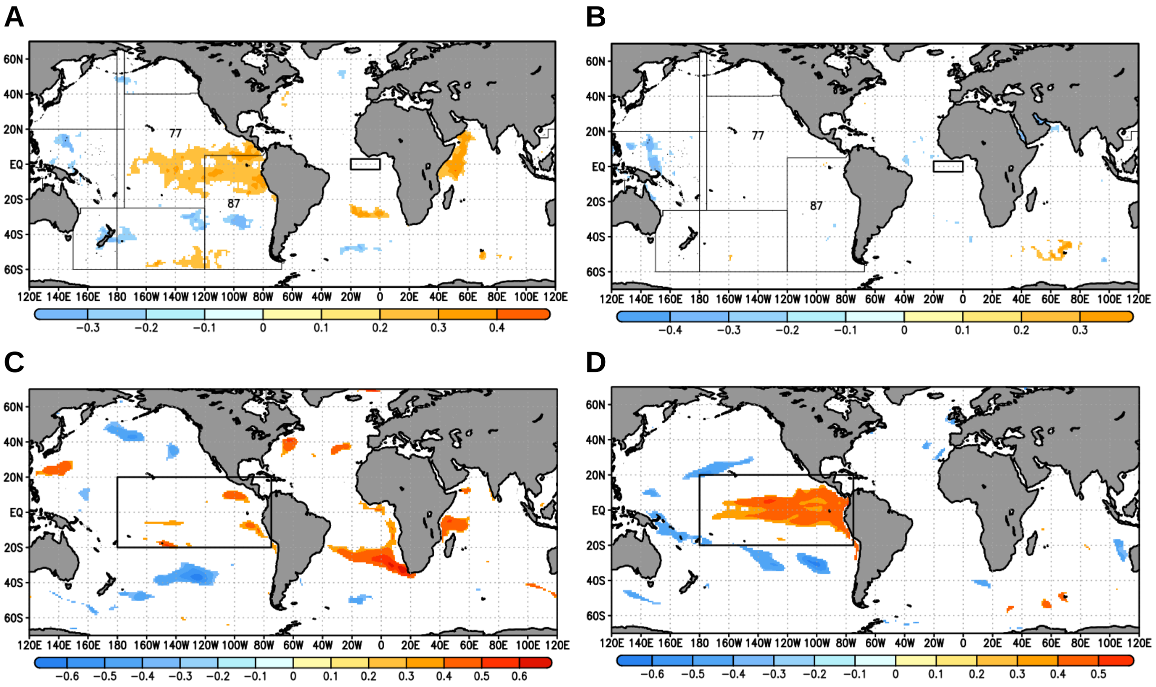


**Fig. S1.** Correlation between Pacific total catch and global SSTs. (*A*) Correlation between interannual Sea Around Us Project historical catch (FAO aggregated 77 & 87 areas) anomalies and second previous calendar year SST anomalies from HadSST (shadings; p-value<0.05). Period 1950-2014. Box for Atl3 SST index calculation in black rectangle. (*B*) Same as (*A*) but for correlation with third previous calendar year SST anomalies. (*C*) Correlation between interannual tropical Pacific averaged (black rectangle) simulated historical total catch anomalies from EcoOcean (EcoG-Fis simulation) and second previous calendar year SST anomalies from GFDL-COBALT (shadings; p-value<0.05). Period 1971-2004. (*D*) Same as (*B*) but for correlation with third previous calendar year SST anomalies.

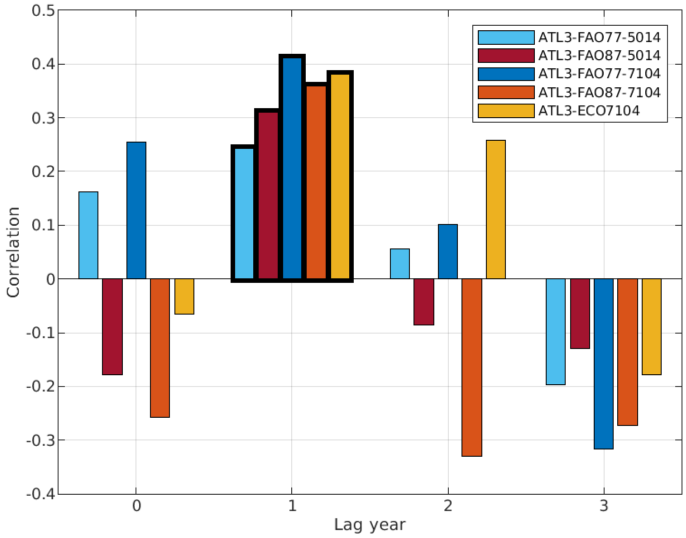


**Fig. S2.** Correlation values (first 4 columns) between Atl3-Had index and year-to-year Sea Around Us Project historical catch data for different FAO major fishing areas (77 & 87), periods (1950-2014 & 1971-2004) and lag years: lag 0 (same calendar year as SST); lag 3 (catch data from 3rd calendar year thereafter). Last column (orange) provides the measure between Atl3-GF index and tropical Pacific averaged [180º-75ºW; 20ºS-20ºN] year-to-year simulated historical total catch anomalies from EcoOcean. Period 1971-2004 (EcoG-Fis simulation). Statistically significant values (95% confidence interval) in thick bar lines.

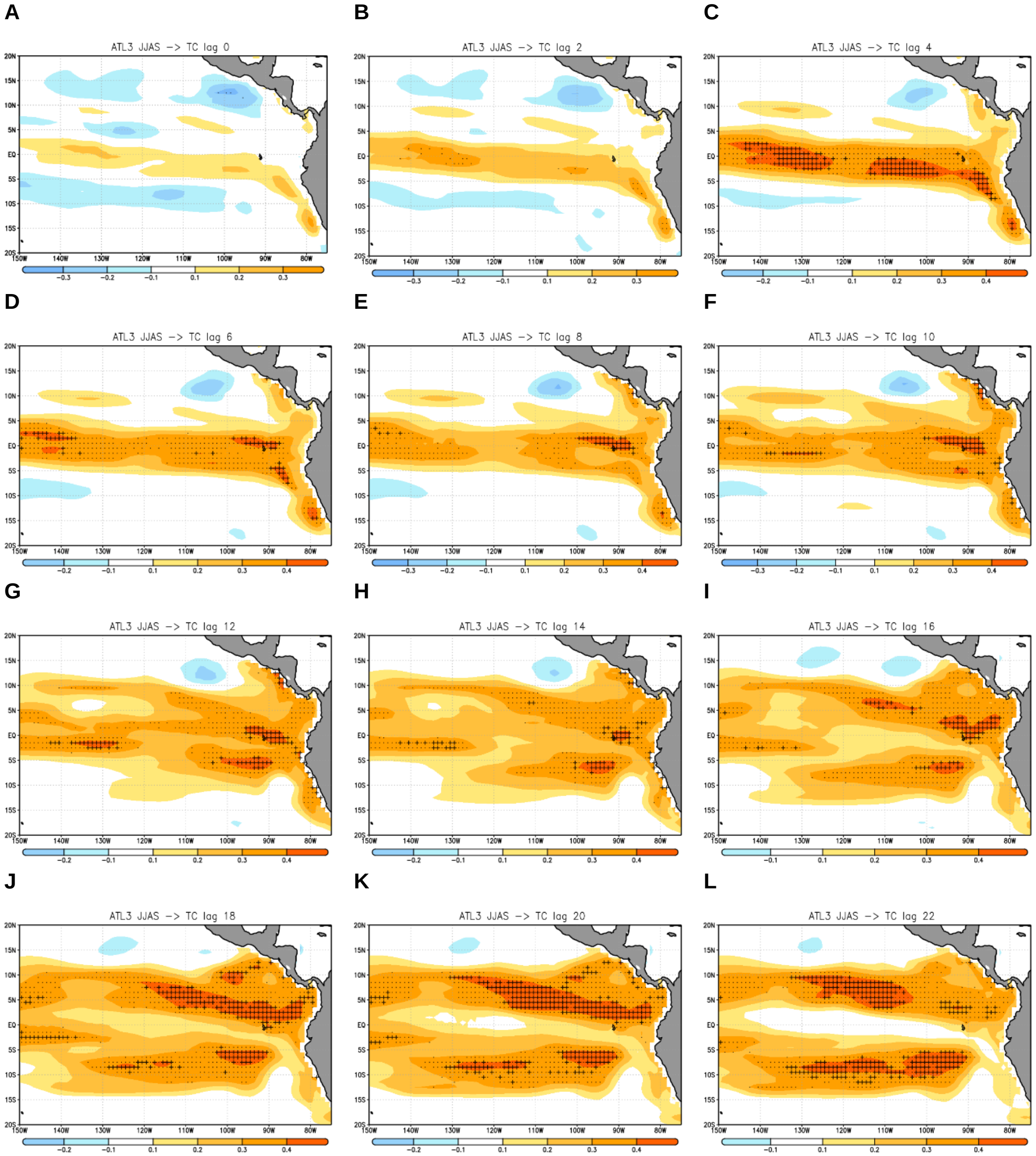


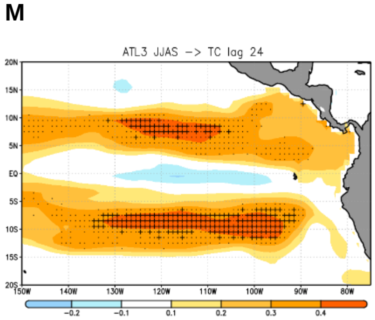


**Fig. S3.** Total catch hindcasts every 2-months. Co-variability maps corresponding to the leading Maximum Covariance Analysis (MCA) modes between equatorial Atlantic lag 0 SST anomalies (predictor; Atl3 region [20ºW-0ºE, 3ºS-3ºN] - black box) and tropical Pacific total catch (tc) anomalies at different monthly lags (predictand; [150ºW-75ºW, 20ºS-20ºN]). (*A-M*) Heterogeneous maps obtained by regression of Us on tc anomalies from monthly lags 0 to 24 (every 2 months). P-values<0.05 in tc-U correlation in stippling. Cross-validated anomaly correlation skill scores calculated from the MCA between hindcasts and observations of tc in crosses (95% confidence interval). Period 1971-2004. EcoG-Fis simulation. See *Methods* section for further information on MCA.

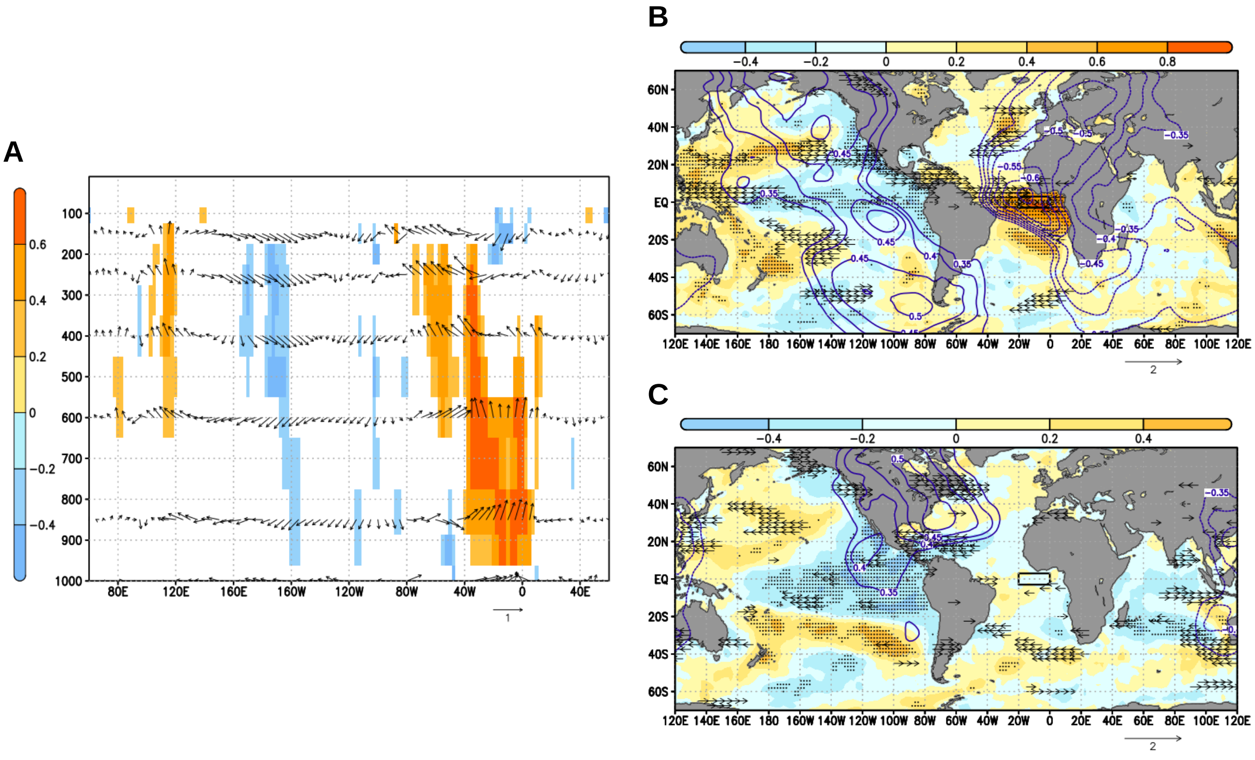


**Fig. S4.** Atlantic-Pacific teleconnection on GFDL-COBALT. (*A*) Correlation between Atl3-GF and meridionally averaged [3ºS-3ºN] NCEP anomalies of zonal wind and omega for lag 0 (JJAS) in arrows ($m\ s^{-1}$/$Pa\ s^{-1}$). Same but for only omega in shadings (p-value<0.05). Omega sign reversed to highlight anomalous air ascent. (*B*) Correlation between Atl3-GF and concomitant SST (shadings in K; p-value<0.05 in stippling; HadISST database), velocity potential at 0.2 sigma level (contours in $10^{-6}\ m^2\ s^{-1}$; p-value<0.05; NCEP database) and zonal wind at 1000 hPa (arrows in $m\ s^{-1}$; p-value<0.05; NCEP database) anomalies. Box for Atl3 calculation in black rectangle. (*C*) Same as (*B*) but for DJFM (lag 5) global anomalies.

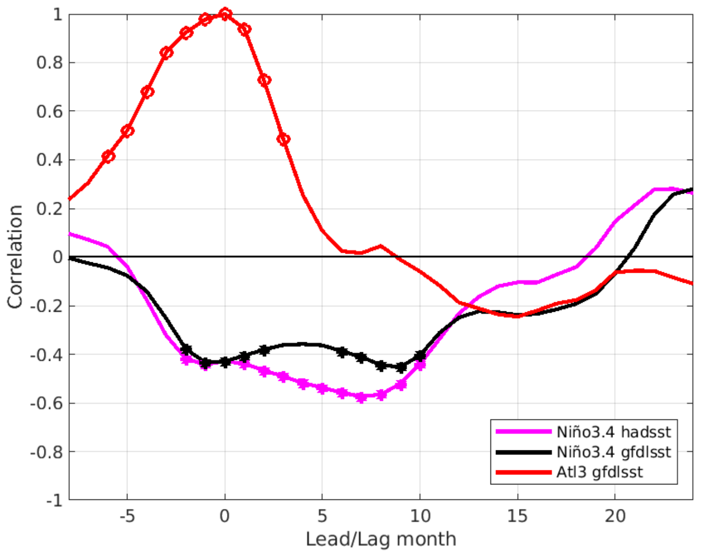


**Fig. S5.** Timing of Atl3 SST anomalies and Atlantic-Pacific teleconnection. Correlation between Atl3-GF and lagged (-6 to 24) Niño3 and Atl3-GF rolling 4-month anomalies of SST (in K; p-value<0.05 in markers). Period 1971-2004.

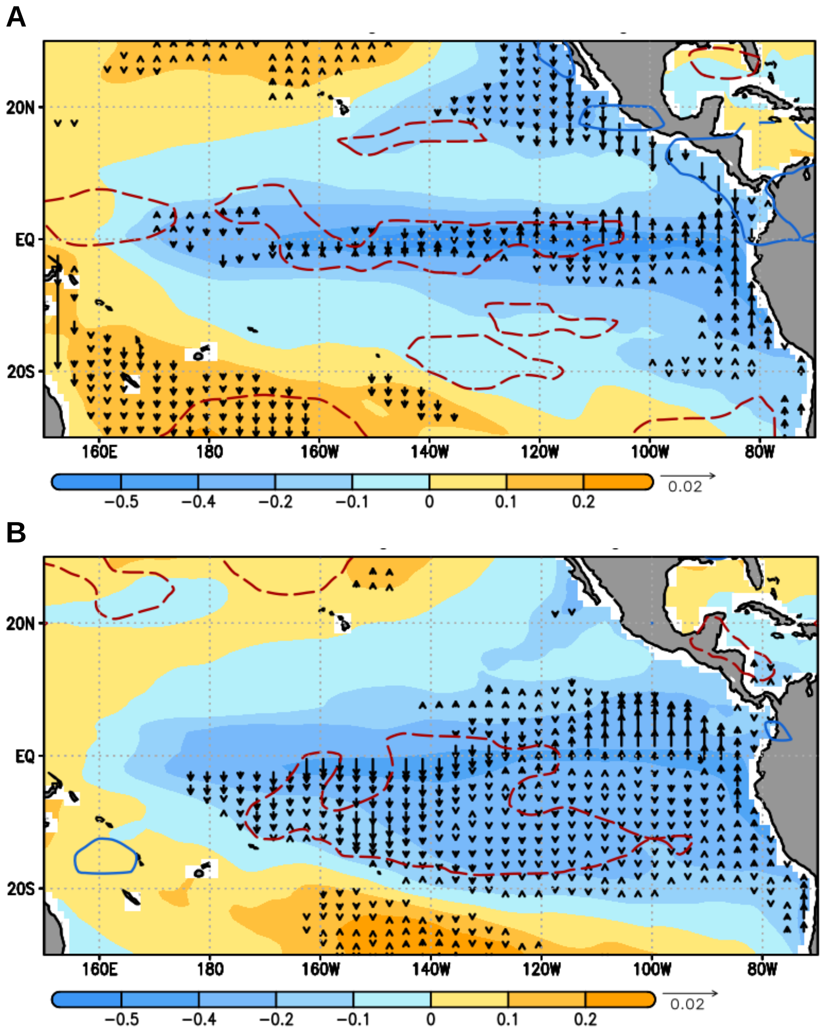


**Fig. S6.** Ekman transport. (*A*) Regression of tropical Pacific anomalous SSTs (shadings in K; GFDL-COBALT) and meridional current at ocean surface (arrows in m $s^{-1}$; only for p-values<0.05 in correlation; GFDL-COBALT) on Atl3-GF. Statistically significant areas for correlation between Alt3-GF and zonal wind at 1000 hPa (m s-1; NCEP) in contours. Lags -1 to +8 (MJJASONDF), 1971-2004. (*B*) Same as (*A*) but for lags +6 to +15 (DJFMAMMJA).

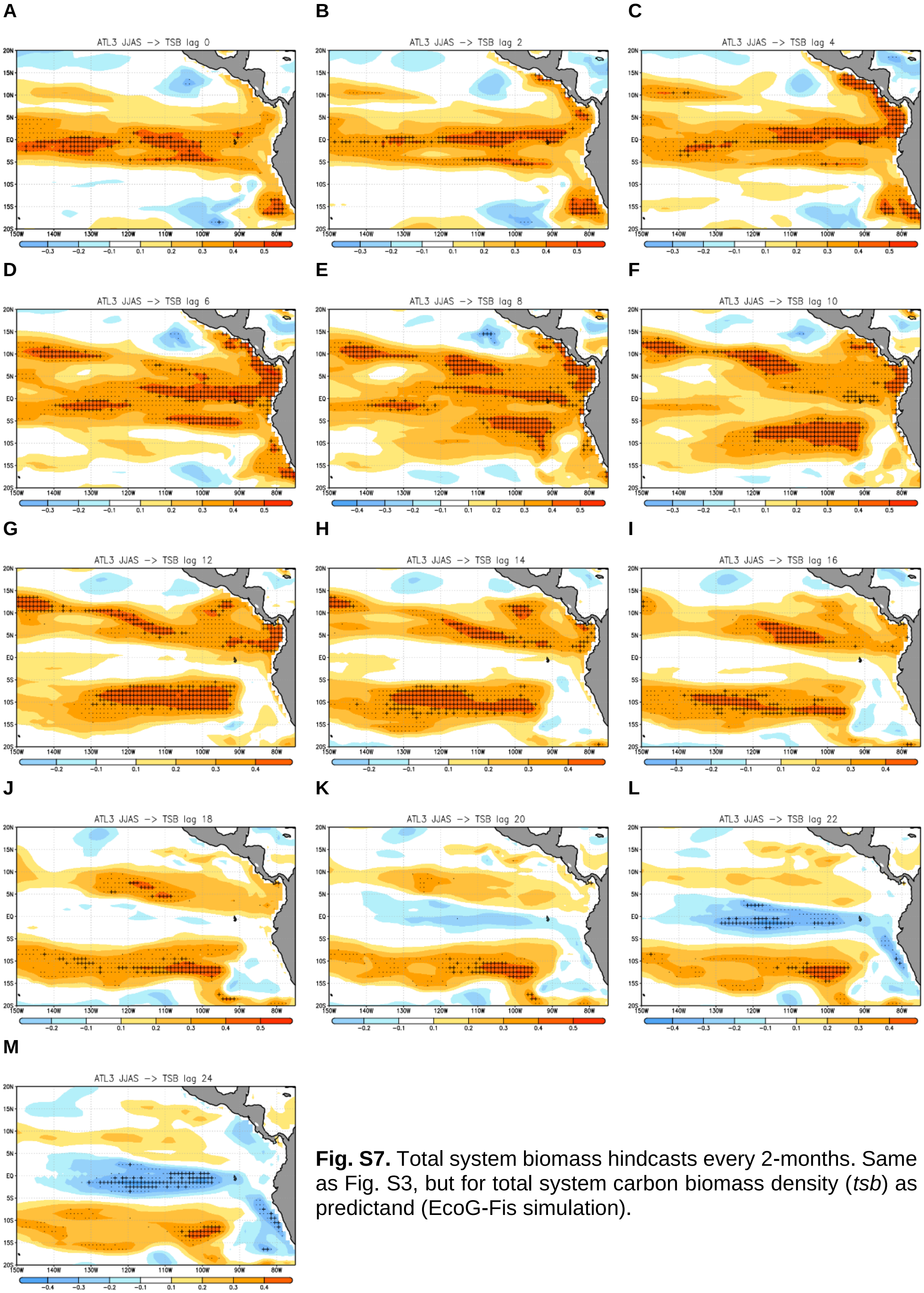


**Fig. S7.** Total system biomass hindcasts every 2-months. Same as Fig. S3, but for total system carbon biomass density (*tsb*) as predictand (EcoG-Fis simulation).

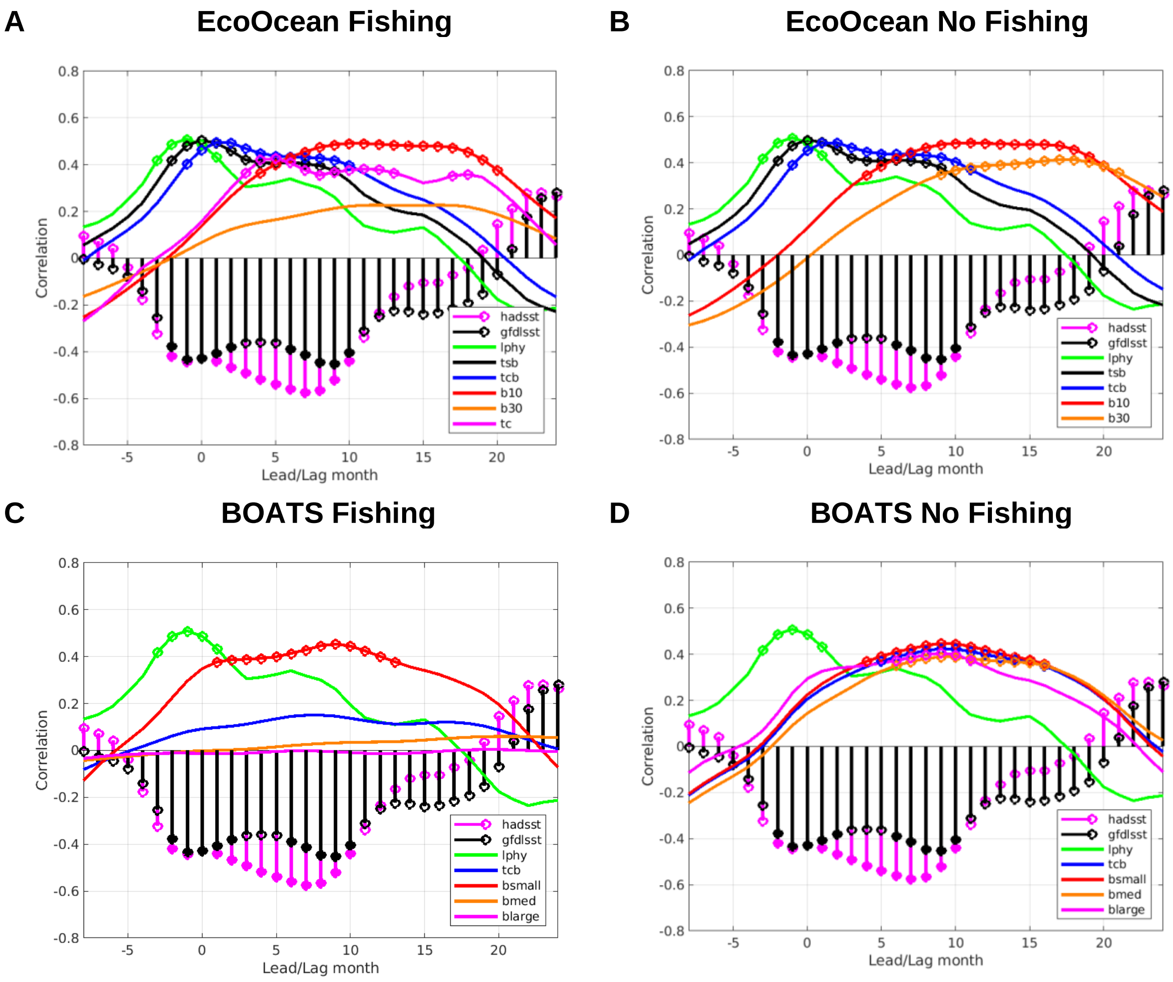


**Fig. S8.** Impact of model setup and fisheries on climate signal propagation I. (*A*) Correlation between Atl3-GF and lagged (-6 to 24) Niño3 rolling 4-month anomalies of SST, large phytoplankton production *(lphy),* total system carbon biomass density *(tsb),* total consumer carbon biomass density *(tcb),* carbon biomass density of consumers greater than 10 cm *(b10)* and 30 cm *(b30*) and total catch at sea *(tc)*. P-value<0.05 in markers (see legend). Period 1971-2004. EcoG-Fis simulation (EcoOcean/Fishing). (*B*) Same as (*A*) but for EcoG simulation (EcoOcean/No-Fishing). (*C-D*) Same as (*A-B*) but for BoaG-Fis (BOATS/Fishing) and BoaG (BOATS/No-Fishing) simulations. Note that *b10* and *b30* are split into *bsmall*, *bmed* and *blarge* in BOATS (cf. Table S1).

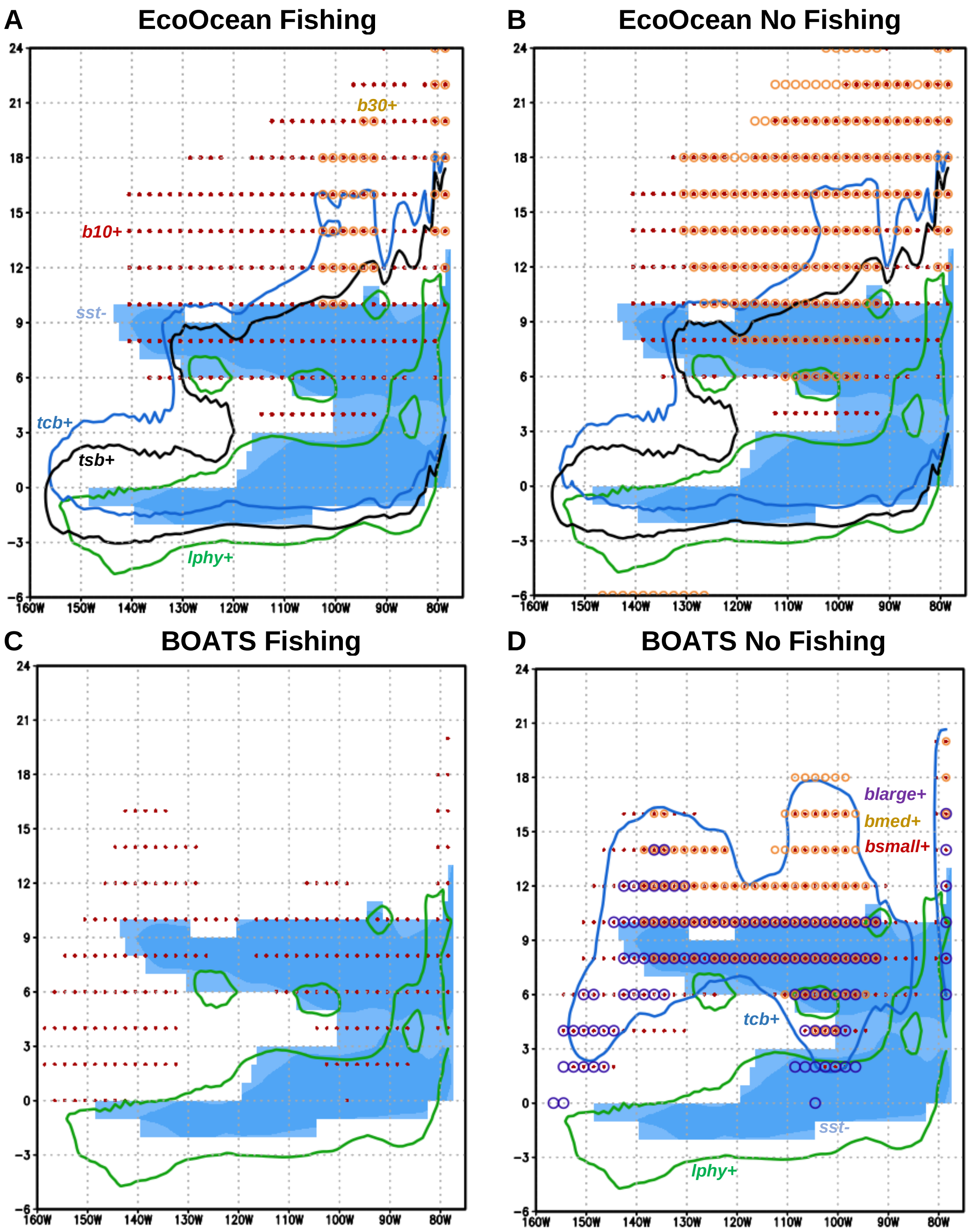


**Fig. S9.** Impact of model setup and fisheries on climate signal propagation II. (*A*) Regression of meridionally averaged [5ºS-5ºN] rolling 4-month anomalies of SST (shadings; in K) on Atl3-GF index, and correlation of meridionally averaged large phytoplankton production (*lphy*; green contours), total system carbon biomass density (*tsb*; black contours), total consumer carbon biomass density (*tcb*; blue contours) and carbon biomass density of consumers greater than 10 cm (*b10*; red dots) and 30 cm (*b30*; orange circles) with the same index. Only p-values<0.05 for positive correlations are shown, except for SST (see text labels). Lags -6 to 24. Period 1971-2004. EcoG-Fis simulation (EcoOcean/Fishing). (*B*) Same as (*A*) but for EcoG (EcoOcean/No-Fishing). (*C-D*) Same as (*A-B*) but for BoaG-Fis (BOATS/Fishing) and BoaG (BOATS/No-Fishing) simulations. Note that in BOATS *b10* and *b30* are split into biomass density of small (*bsmall*; red dots), medium (*bmed;* orange circles) and large size *(blarge;* magenta circles*)* consumers and *tsb* (black curve) is not available (cf. Table S1).

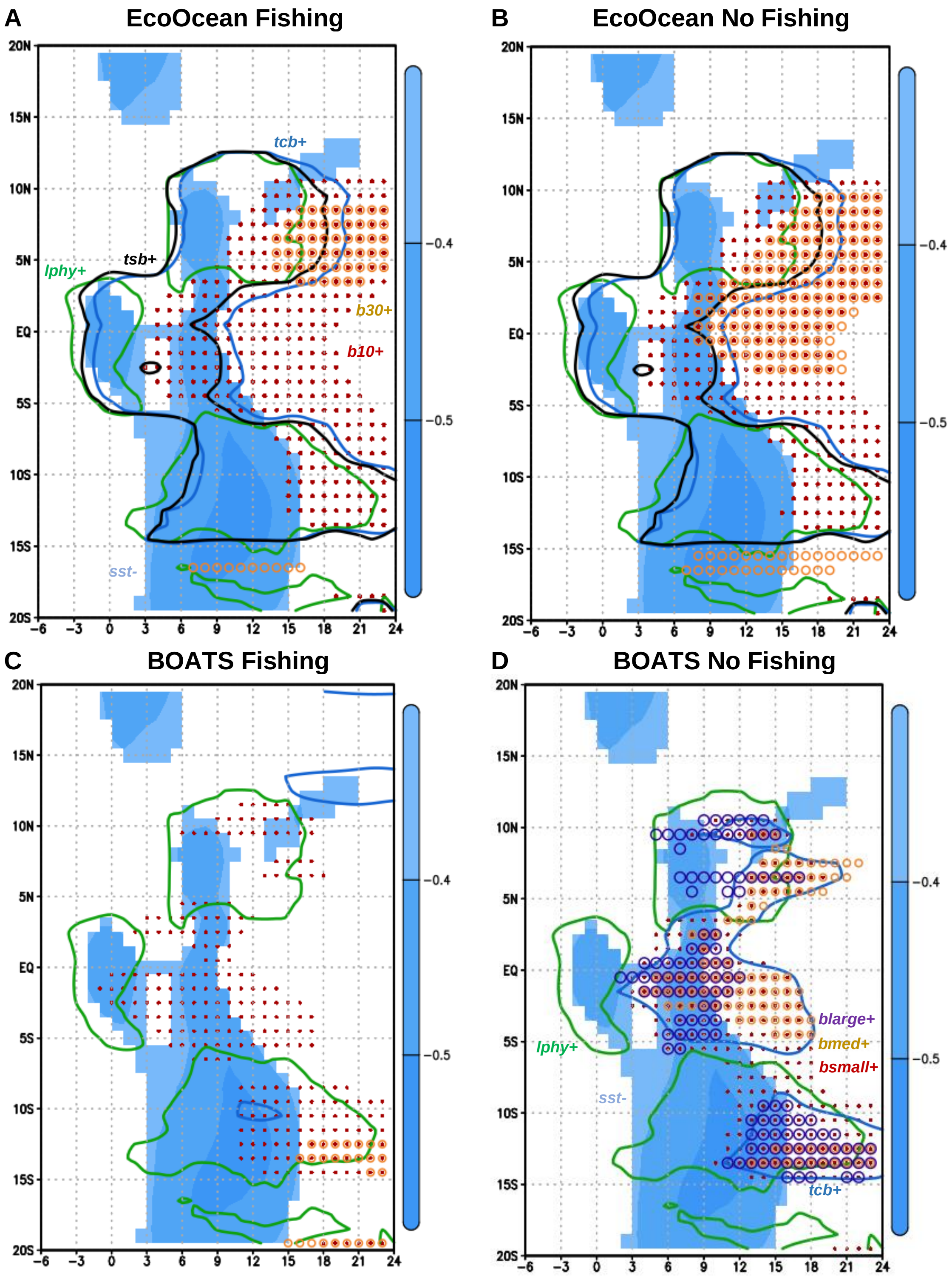


**Fig. S10.** Impact of model setup and fisheries on climate signal propagation III. Same as Fig. S9, but for zonally averaged [150ºW-75ºW] anomalies.

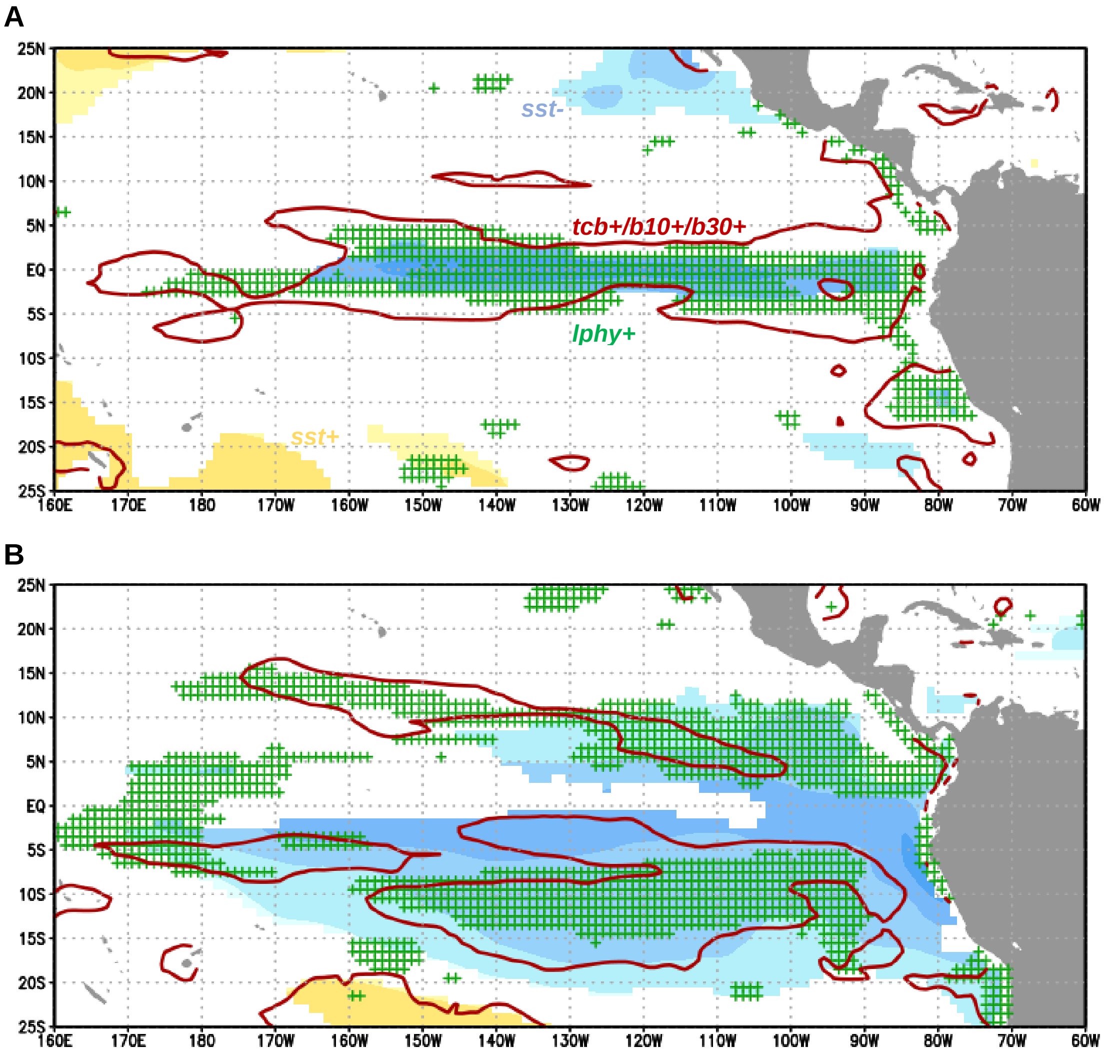


**Fig. S11.** Atlantic-Pacific teleconnection impact on Macroecological. (*A*) Regression of tropical Pacific SST annual anomalies (shadings; in K) on Atl3-GF index on same calendar year, and correlation of large phytoplankton production (*lphy*; green crosses) and total system carbon biomass density/carbon biomass density of consumers greater than 10 cm/carbon biomass density of consumers greater than 30 cm (*tsb/b10/b30*; red contours) annual anomalies with the same index. Only p-values<0.05 for positive correlations are shown, except for SST (see text labels). (*B*) Same as (*A*) but for variables from calendar year after. Period 1971-2004. GFDL-COBALT and MacG simulations (Macroecological/No fishing).

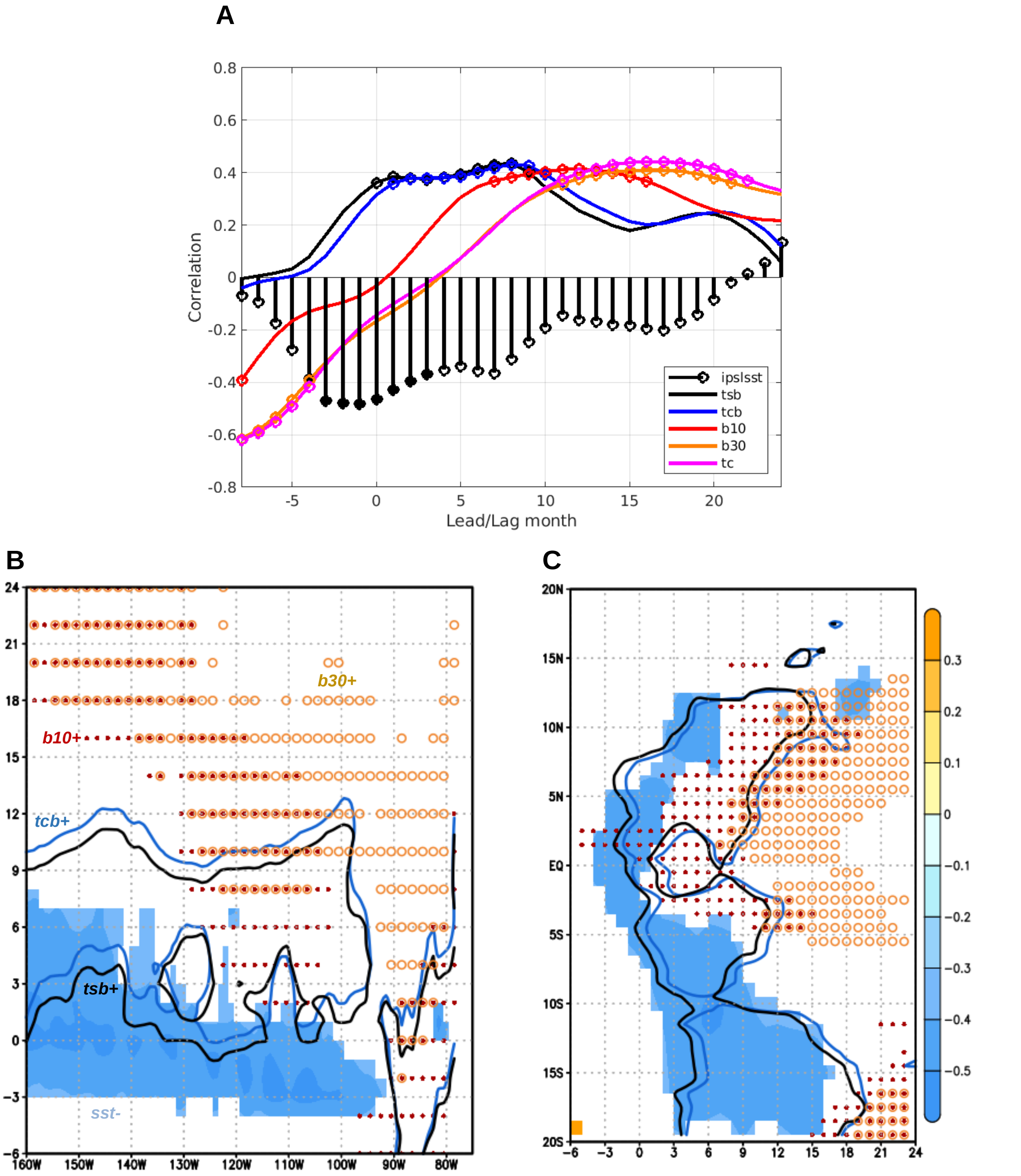


**Fig. S12.** Predictability of tropical Pacific living marine resources under climate change. (*A*) Correlation between Atl3 JJAS SST from IPSL-CM5A-LR and lagged (-6 to 24) Niño3 rolling 4-month anomalies of SST, total system carbon biomass density (*tsb*), total consumer carbon biomass density (*tcb*), carbon biomass density of consumers greater than 10 cm *(b10)* and 30 cm *(b30*) and total catch at sea *(tc)*. P-values<0.05 in markers. (*B*) Regression of meridionally averaged [5ºS-5ºN] rolling 4-month anomalies of SST (shadings; in K) on Atl3 JJAS SST IPSL-CM5A-LR, and correlation of meridionally averaged *tcb* (blue contours), *b10* (red dots) and *b30* (orange circles) with the same index. Only p-values<0.05 for positive correlations are shown, except for SST (see text labels). Lags -6 to 24. (*C*) Same as (*B*) but for zonally averaged anomalies [150ºW-75ºW]. Period 2021-2054. EcoI8.5-Fis simulation (EcoOcean/Fishing).
